# Elevated plasma cortisol associated with larger ventricles and smaller hippocampal volumes – a study in 2 independent elderly cohorts

**DOI:** 10.1101/2020.05.03.074823

**Authors:** Priya Rajagopalan, Kwangsik Nho, Shannon L. Risacher, Neda Jahanshad, Christina P. Boyle, Andrew J. Saykin, Paul M. Thompson

**Author notes:** Senior authors AJS and PMT jointly supervised the work. Parnassus avenue, Room L352, San Francisco, CA 94143. Corresponding author: Priya Rajagopalan (,).

## Abstract

Cortisol is considered the most fundamental stress hormone and is elevated in stress and multiple neuropsychiatric conditions. Prior studies have shown associations of plasma cortisol levels with total cerebral and hippocampal volumes and less consistently with the amygdala. Here, we extend our hypothesis to test associations of plasma cortisol with 1) ventricular 2) hippocampal and 3) amygdalar volumes, in two independent elderly cohorts across a broad cognitive spectrum ranging from normal cognition to Alzheimer’s disease.

We demonstrate elevated cortisol to be associated with larger lateral ventricular volumes and smaller hippocampal volumes, predominantly in the right cerebral hemisphere, regardless of age, sex or cognitive status. We noted a non-significant trend of smaller amygdalar volumes with elevated cortisol.

Our findings support smaller brain parenchyma volumes seen with elevated cortisol and may encourage effective strategies reducing cortisol and stress. They may also serve as imaging biomarkers for assessing therapeutic benefits of stress and cortisol lowering interventions aiming to halt or reverse the brain volume alterations and theoretically improve cognition and quality of life.

**Highlights:**

- Elevated cortisol associated with larger ventricular volumes and smaller hippocampal volumes
- Associations are predominantly noted in the right cerebral hemisphere.
- Similar non-significant trends noted in amygdalar volumes
- Cortisol and stress reducing strategies may halt brain changes and improve quality of life
- Imaging biomarkers may help assess efficacy of cortisol-lowering therapeutic interventions

## 1. Introduction

Stress is an inevitable challenge in everyday life. Typically, the human body responds to stressful stimuli by triggering the release of cortisol into the bloodstream through activation of the hypothalamic-pituitary-adrenal (HPA) axis, which facilitates restoration of homeostasis. Cortisol, a corticosteroid hormone, is considered a reliable indicator of HPA activity and is the most fundamental stress hormone. Excessive and sustained production of cortisol can cause malfunctioning of the HPA feedback loop, leading to symptoms of physiological and psychological distress (Chrousos and Gold, 1992; Le Fevre et al., 2003), all of which can increase morbidity and mortality (Pratt, 2009).

While cortisol levels increase with physiological aging (Ferrari et al., 1995; Ferrari et al., 2004), multiple neuropsychiatric conditions demonstrate elevated cortisol levels irrespective of age. For example, cortisol levels are higher than normal in conditions such as schizophrenia, bipolar disorder (Gallagher et al., 2007; Ryan et al., 2004), major depression (Linkowski et al., 1987; Plotsky et al., 1995) and Alzheimer’s disease (Davis et al., 1986; Martignoni et al., 1990). Within these conditions, cortisol levels are higher in those with a greater degree of cognitive impairment (Hinterberger et al., 2013; Zvěřová et al., 2013).

The limbic system of the brain, including the hippocampus and amygdala, has abundant corticosteroid receptors, making it a principal target for increased vulnerability to prolonged cortisol exposure. There are two main kinds of corticosteroid receptors expressed in the brain, most notably in the hippocampus, called type I mineralocorticoid receptors with permissive effects and type II glucocorticoid receptors with suppressive effects (de Kloet et al., 2005; Sapolsky, 1996). The hippocampus is involved in the consolidation and retrieval of declarative memory, whereas the amygdala supports affective and emotional aspects of cognition (Pruessner et al., 2010). These are both important structures for HPA axis regulation. The hippocampus is primarily inhibitory and exerts a negative feedback on the HPA axis after termination of a stressful stimulus, whereas the amygdala is predominantly excitatory in nature.

Neuroimaging studies have found elevated plasma cortisol levels to be associated with smaller hippocampi in AD (De Leon et al., 1988; Huang et al., 2009) and in cognitively healthy older adults (Lupien et al., 1998). Plasma cortisol levels have also been shown to be associated with smaller total cerebral brain volumes in healthy middle aged women, but not in men of middle-age (Echouffo-Tcheugui et al., 2018) or older ages (MacLullich et al., 2005). Additionally, evening but not morning, salivary cortisol levels were shown to be associated with smaller total brain, gray and white matter volumes in older adult participants without dementia (Geerlings et al., 2015).

Both hippocampi and amygdala demonstrate increased glucocorticoid receptors in hypercortisolemic neuropsychiatric conditions such as major depressive disorder (Wang et al., 2012; Wang et al., 2014) and brain volume reductions have been reported in early onset depression (Janssen et al., 2004; Schmaal et al., 2016). While stress and hypercortisolemic neuropsychiatric diseases have consistently shown smaller hippocampi (Pruessner et al., 2010; Sapolsky, 1996; Swaab et al., 2005), the volumetric associations for the amygdala however have been inconsistent (Kronenberg et al., 2009; Malykhin et al., 2018; Mervaala et al., 2000; Mitra et al., 2005; Pruessner et al., 2010).

In this study, we investigated the association of plasma cortisol levels with region of interest (ROI)-based subcortical volumes in two independent cohorts of older adults across a broad cognitive spectrum, including normal cognition (CN), mild cognitive impairment (MCI), and Alzheimer’s disease (AD). We focused on established neuroimaging markers for neuropsychiatric disorders such as AD, including lateral ventricular, hippocampal and amygdalar volumes (Hendren et al., 2000).

## 2. Materials and methods

The study was conducted according to the Good Clinical Practice guidelines, the Declaration of Helsinki, and US 21 CFR Part 50 — Protection of Human Participants and Part 56 — Institutional Review Boards in the two cohorts, namely the Alzheimer’s Disease Neuroimaging Initiative (ADNI) and the Indiana Memory and Aging Study (IMAS). All study participants gave written informed consent.

### 2.1 ADNI

Data used in preparing this article were obtained from the Alzheimer’s Disease Neuroimaging Initiative (ADNI) database (http://adni.loni.usc.edu/). ADNI is a multisite study that includes older adults across a spectrum of cognitive impairment; each participant was assessed with at least one 1.5 tesla anatomical brain MRI scan, blood sampling, and cognitive and clinical testing at one of 58 sites across North America. Inclusion and exclusion criteria for ADNI are detailed online at http://adni.loni.usc.edu (Mueller et al., 2005). The present analysis was carried out in a total of 487 non-Hispanic White participants (average age: 75.13 +/− 0.3 years; 301 males and 186 females; 107 with AD, 332 with MCI and 48 CN) with brain MRI and plasma cortisol data.

### 2.2 IMAS

The Indiana Memory and Aging Study (IMAS) is a regional cohort, based in Indianapolis, Indiana, which includes older adults across a broad cognitive spectrum. The selection criteria and characterization have been described previously (Kim et al., 2013; Risacher et al., 2013; Saykin et al., 2006). All participants underwent a 3D T1-weighted MPRAGE brain scan on a Siemens TIM Trio 3T scanner, as well as collection of blood samples, and extensive cognitive and clinical testing. The analysis was performed in a total of 54 non-Hispanic White participants (average age: 72.57 +/− 0.98 years; 19 males and 35 females; 6 with AD, 21 with MCI and 27 CN) who had plasma cortisol and brain MRI data.

### 2.3 MRI acquisition

The T1-weighted 1.5 T structural brain MPRAGE scans were acquired at multiple ADNI sites with a standardized protocol (Jack et al., 2010). The T1-weighted 3T structural brain MRI scans were acquired for the IMAS participants following the ADNI protocol.

### 2.4 Regional brain volumes

For the analysis, total intracranial volumes, left and right lateral ventricular, hippocampal, and amygdalar volumes (in mm^3^) were extracted from MRI scans for both ADNI and IMAS participants using the FreeSurfer image analysis suite, version 5.1 (http://surfer.nmr.mgh.harvard.edu/; (Fischl et al., 2002)).

### 2.5 Morning plasma cortisol levels

Plasma cortisol exhibits a circadian rhythmicity with highest levels being secreted in the morning (Chida and Steptoe, 2009). In ADNI, plasma cortisol levels were obtained from morning blood samples collected at the time of MRI scan (http://www.adni-info.org/Scientists/Pdfs/14-Biomarker_Sample_Collection_Processing_and_Shipment.pdf) using the ‘Human Discovery Multi-Analyte Profile’ platform by Myriad Rules-Based Medicine (RBM, www.rulesbasedmedicine.com, Austin, TX). The quantification method is described in the document ‘Biomarkers Consortium ADNI Plasma Targeted Proteomics Project – Data Primer’ (http://adni.loni.usc.edu/wp-content/uploads/2010/11/BC_Plasma_Proteomics_Data_Primer.pdf). Plasma cortisol assessment for the IMAS mirrored the ADNI protocol and has been described previously (Kim et al., 2013).

Theoretically, altered HPA axis activity with artificially low cortisol levels may result from including steroid containing oral medications and intra-articular injections (Habib, 2009; Hengge et al., 2006; Lipworth, 1999). Therefore, after reviewing the medication history in detail, 30 of the 517 (5.8%) participants from ADNI with steroid usage (oral, injection and inhalational route of usage) were identified and excluded from the final analysis which was performed in 487 participants. Of note, the inclusion or exclusion of these participants in the reported analyses did not alter the pattern of significant results (data not presented here). Similarly, 4 of 58 (6.9%) of the IMAS participants were excluded for usage of steroids and the final analysis was performed in 54 participants.

### 2.6 Statistical analysis

To reduce skewness in the plasma cortisol concentrations, we carried out a logarithmic transformation of the cortisol measure using the natural logarithmic transformation (Figure 1).

**Figure 1.**
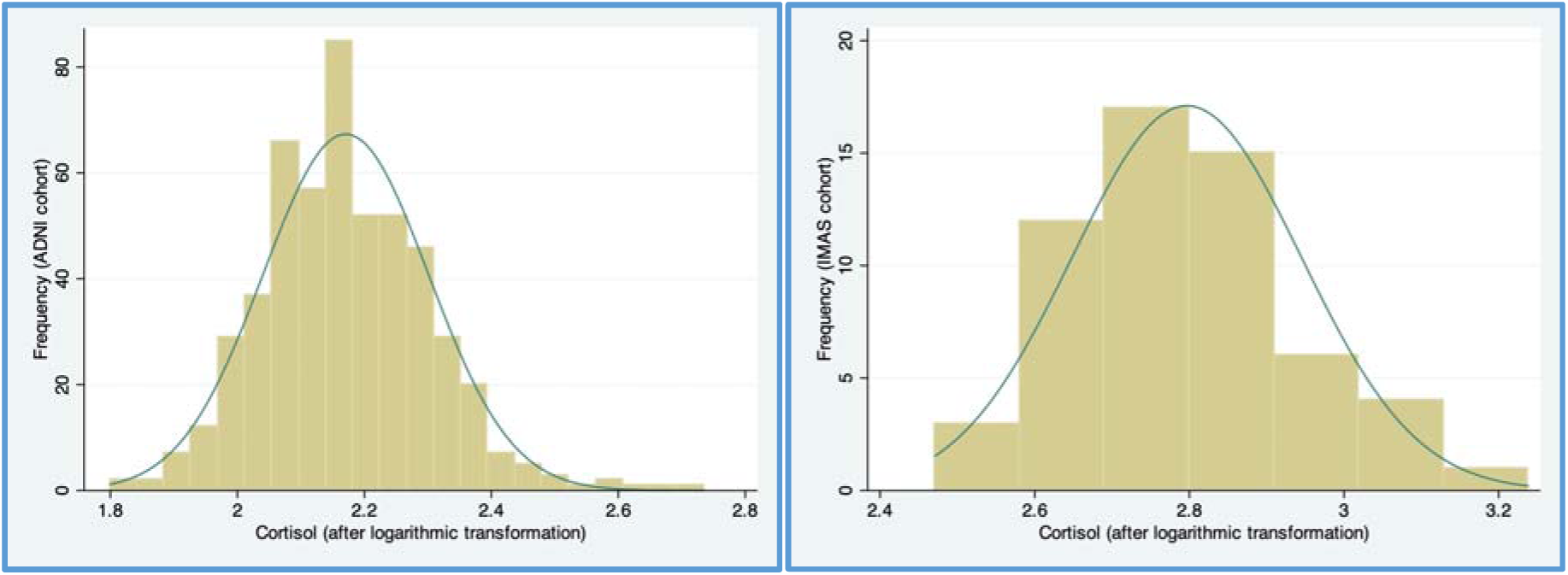
Logarithmic transformation of plasma cortisol measures using the natural logarithm was carried out to reduce skewness of the data in both cohorts.

Using linear regression performed in the *Stata* (StataCorp, 2011, College Station, Texas) software package, we modeled the effect of cortisol measures (Equation 1) on left and right lateral ventricular, hippocampal and amygdalar volumes (in mm^3^), in both the ADNI and IMAS cohorts. The association of plasma cortisol levels with brain subcortical volumes was first assessed in ADNI (Table 1a) and tested for replication in IMAS (Table 1b). An additional *post-hoc* mega-analysis was carried out by combining both ADNI and IMAS participants (Table 1c).

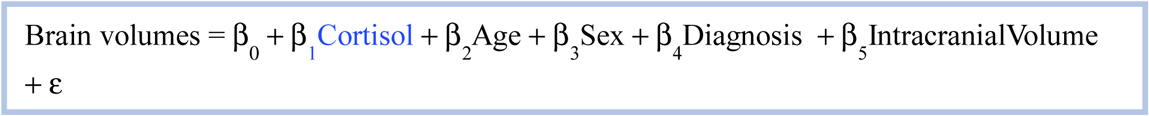

**Table 1.**
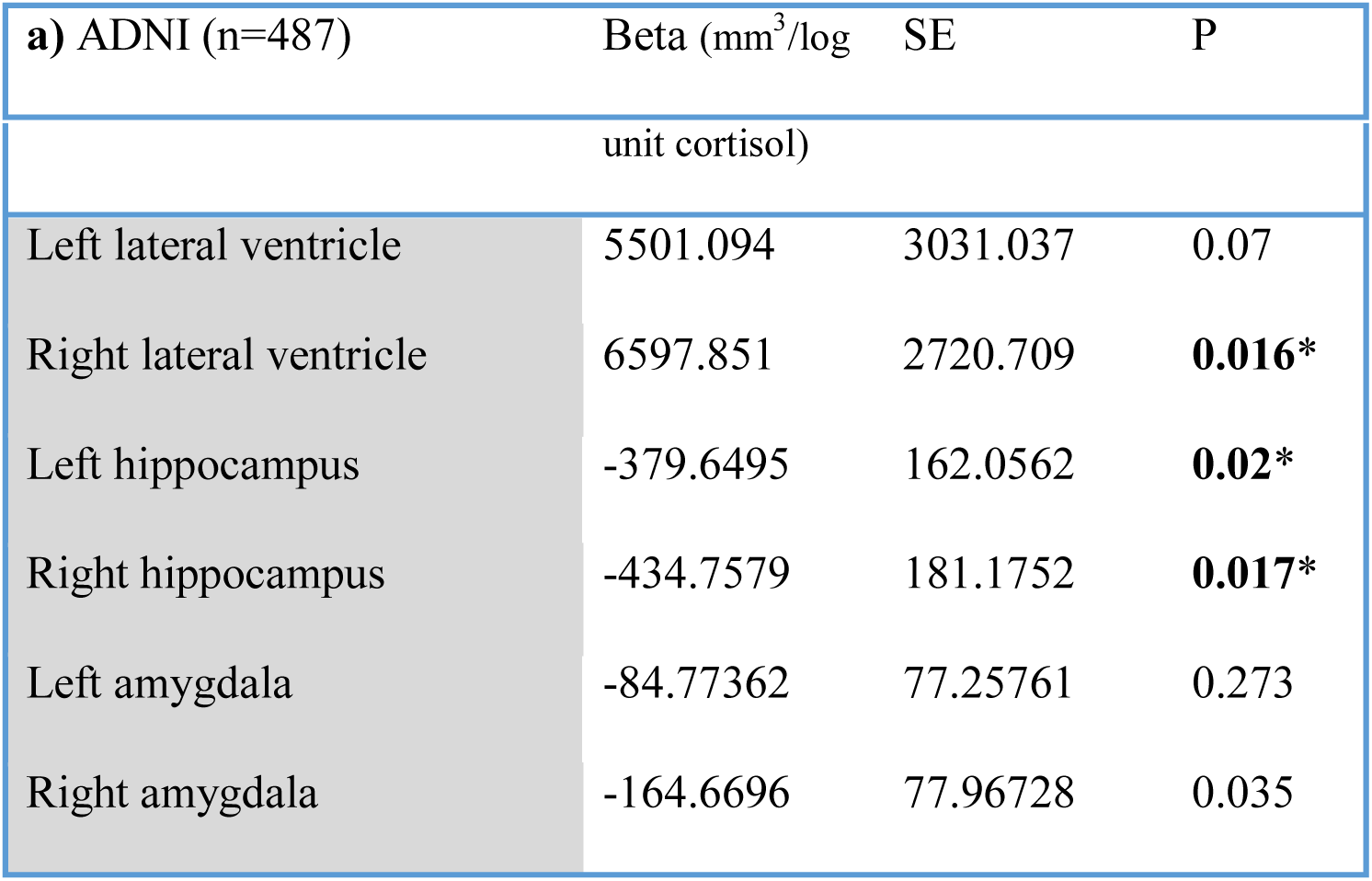
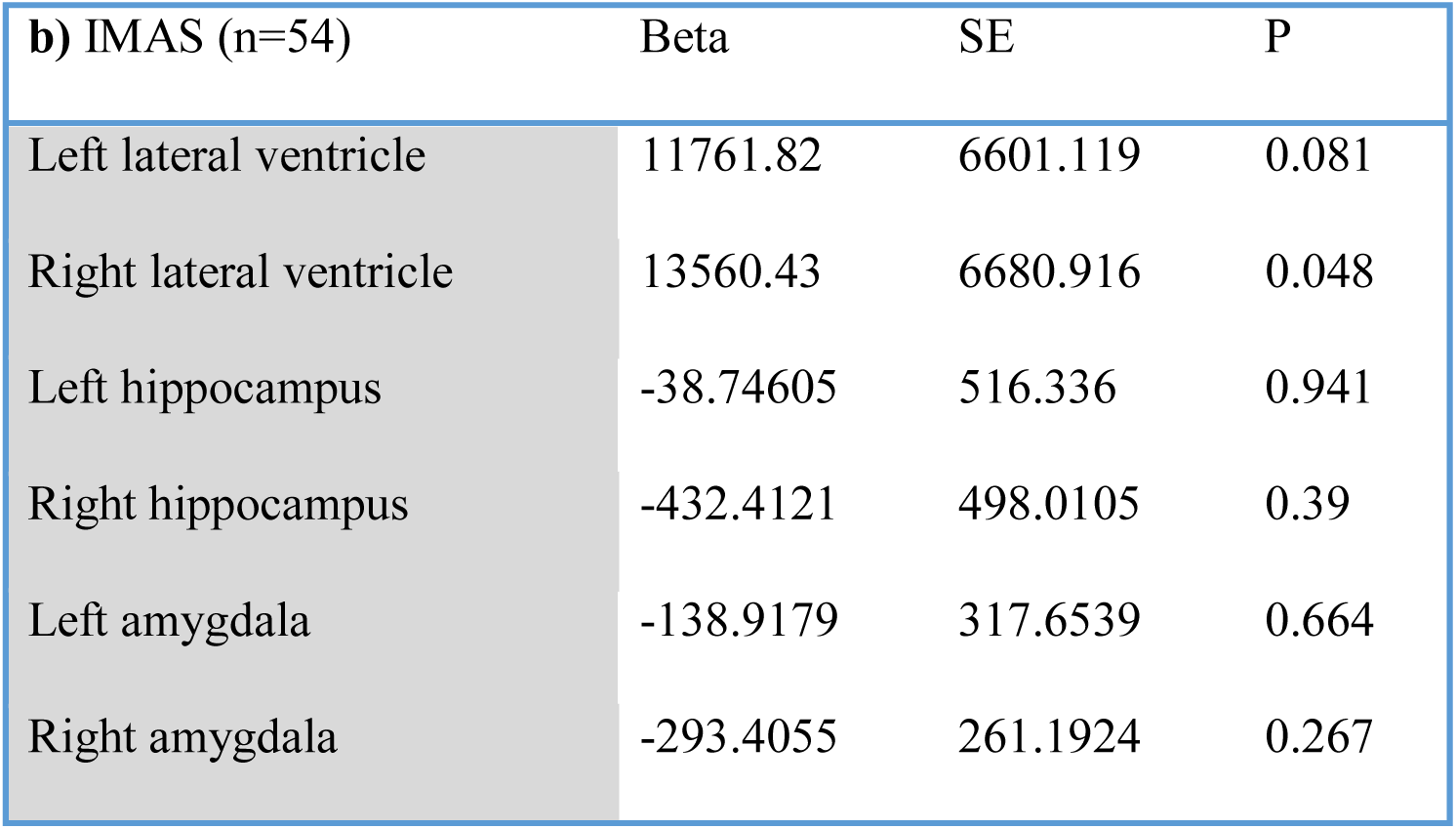
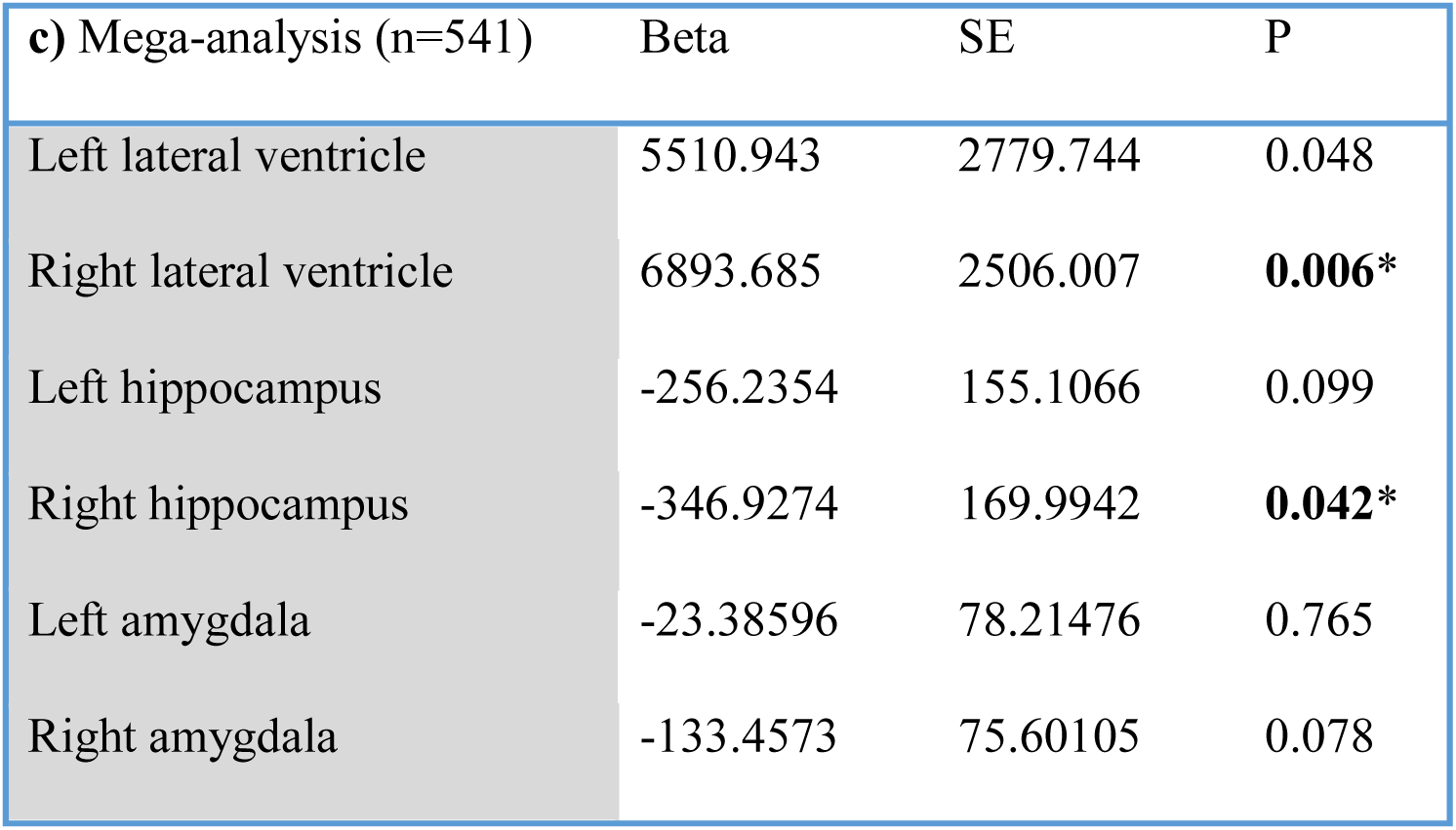
**(a)** Linear regression analysis demonstrates a significant association of cortisol levels with ventricular and hippocampal volumes in older adults in the ADNI cohort, based on PC analysis adjusted Bonferroni-corrected P-value of <0.025 considered significant. **(b)** There were non-significant but similar trend of associations noted in the much smaller IMAS cohort. **(c)** A mega-analysis combining both ADNI and IMAS data demonstrated significant association of cortisol levels with ventricular and hippocampal volumes, using a P-value < 0.05. All associations were adjusted for effects of age, sex and diagnosis (AD, MCI, or CN). Significant P-values are denote by *.

Equation 1: Linear regression model used in the study

All associations were adjusted for predictors regarded as potential confounders, including age, sex, total intracranial volume and diagnosis (AD, MCI or CN). Age and sex were known confounders affecting the HPA axis and the HP gonadal axis, with studies demonstrating an accentuated cortisol response in older women (Otte et al., 2005). We adjusted for total intracranial volume to eliminate confounding effects from variation in head size. Covariates include diagnostic status of AD, MCI or CN to eliminate the known association of hypercortisolism with AD (Davis et al., 1986; Martignoni et al., 1990) and cognition (Echouffo-Tcheugui et al., 2018).

As the six imaging phenotypes (left and right ventricular, left and right hippocampal and left and right amygdalar volumes) strongly correlate with each other, we applied principal component analysis to estimate the number of independent principal components (PC) and to adjust for multiple testing with Bonferroni correction. Two PCs were found to be significant (with an eigenvalue > 1) and these components explained 87% of the variation of the six regions of interest. Thus, an association with p-value < 0.025 (0.05/2) was considered as significant. Volumetric regression analysis of cortisol histogram plots and regional brain volumes and were created using *Stata.*

## 3. Results

Of the 487 ADNI participants (average age: 75.13 +/− 0.3 years; 301 males and 186 females), 107 were diagnosed with AD, 332 with MCI and 48 were CN. Of the 54 IMAS participants (mean age: 72.57 +/− 0.98 years; 19 males and 35 females), 6 were diagnosed with AD, 21 with MCI and 27 were CN.

The mean and standard deviation of logarithmically transformed cortisol levels in the ADNI cohort were 2.17+/− 0.13. logarithmic units (2.20 +/− 0.11 in AD, 2.16 +/− 0.13 in MCI and 2.17 +/− 0.15 in CN) whereas in IMAS cohort were 2.80 +/− 0.15 logarithmic units (2.20 +/− 0.11 in AD, 2.16 +/− 0.13 in MCI and 2.17 +/− 0.15 in CN). Histogram plots with overlaid normal curves for cortisol levels are demonstrated for both cohorts in Figure 1.

Higher plasma cortisol measures were associated with significantly larger right lateral ventricular volumes (*p=0.016*) in ADNI (Table 1a & Figures 2b) and in a mega-analysis combining ADNI and IMAS participants (*p=0.006*) (Table 1c), with similar non-significant trends noted in the smaller IMAS cohort (*p=0.048*) (Table 1b & Figure 3b). Both ADNI and IMAS cohorts demonstrated similar but non-significant trends of larger left ventricular volumes associated with elevated cortisol.

**Figure 2:**
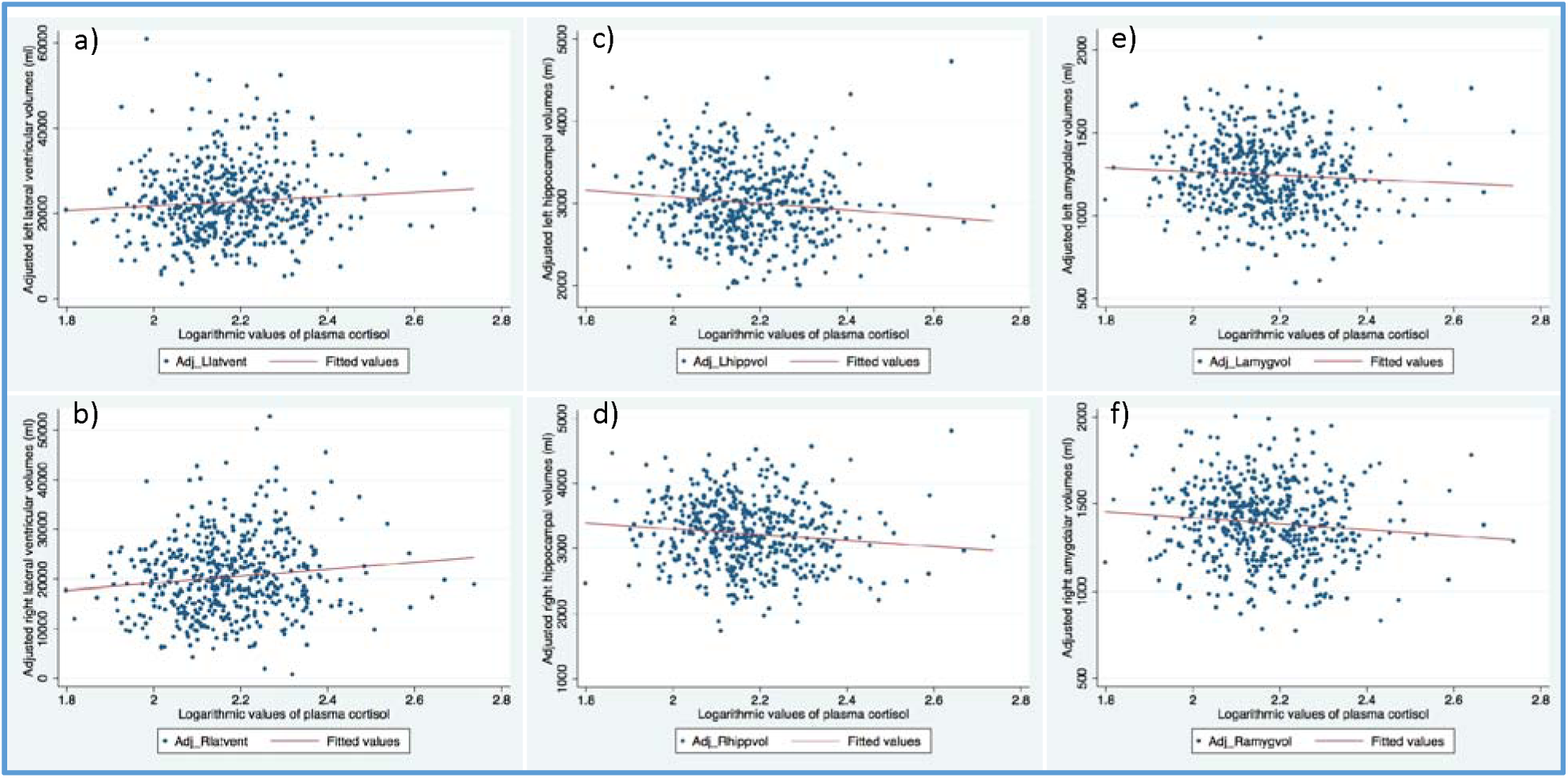
In ADNI, elevated cortisol was associated with greater **(a)** left lateral ventricular (p=0.07) and **(b)** right lateral ventricular (p=0.016*) volumes, as well as lower **(c)** left hippocampal (p=0.02*), **(d)** right hippocampal (p=0.017*), **(e)** left amygdalar (p=0.273) and **(f)** right amygdalar (p=0.035) volumes, after adjusting for effects of age, sex, intracranial volumes and diagnosis (AD, MCI or CN) and using a Bonferroni corrected P < 0.025.

**Figure 3:**
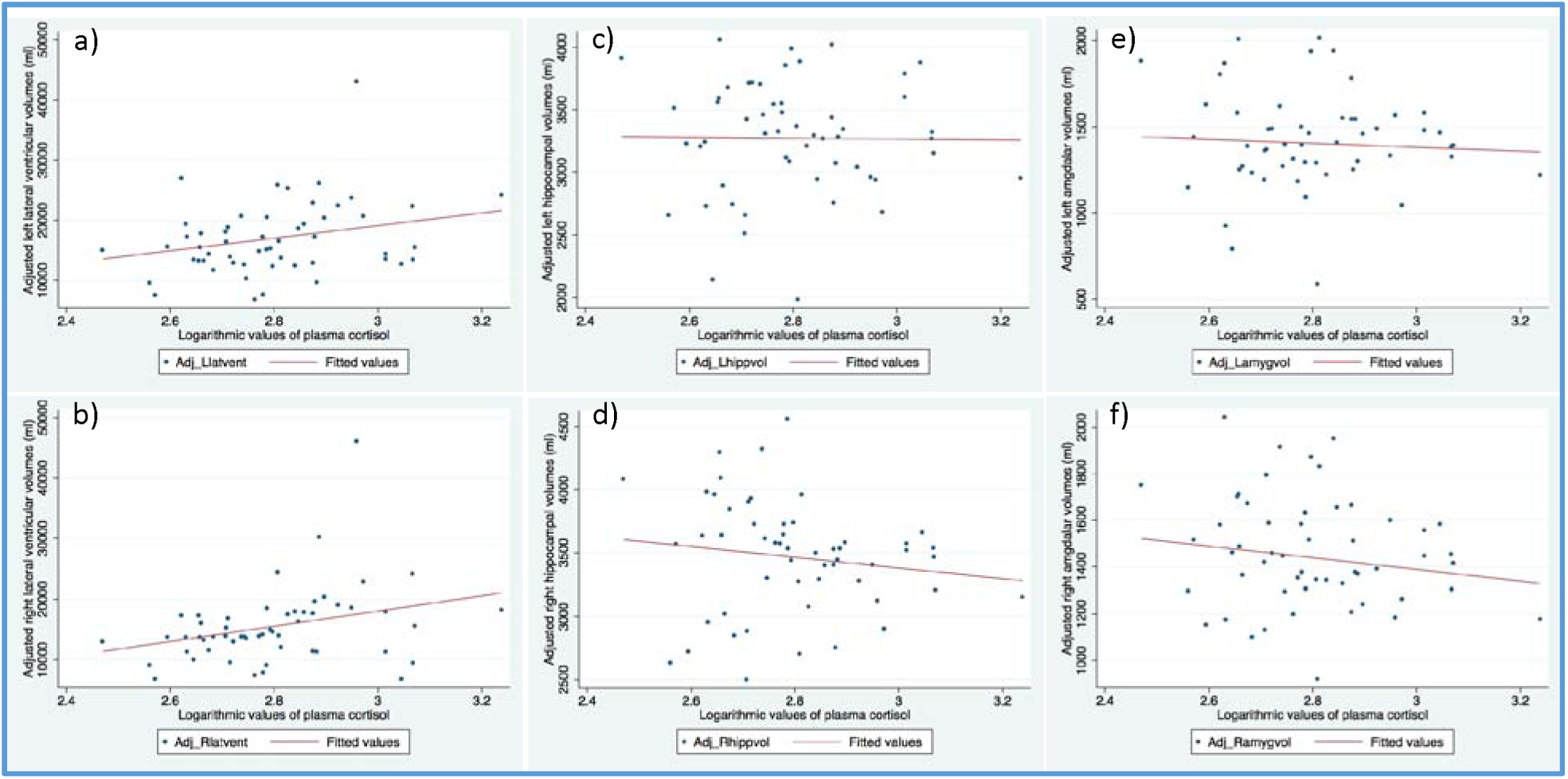
In IMAS, elevated cortisol was associated with non-significant but similar trends of greater **(a)** left lateral ventricular and **(b)** right lateral ventricular volumes, as well as lower **(c)** left hippocampal, **(d)** right hippocampal, **(e)** left amygdalar and **(f)** right amygdalar volumes, after adjusting for effects of age, sex, intracranial volumes and diagnosis (AD, MCI or CN).

In ADNI participants, elevated cortisol levels were associated with significantly smaller right hippocampal (*p=0.02*) and left hippocampal volumes (*p=0.017*) (Table 1a, Figure 2c, **2d)**. In a mega-analysis combining ADNI and IMAS participants right hippocampal volumes continued to be significantly associated (*p=0.042*) (Table 1c), with similar but non-significant trends demonstrated in the smaller IMAS cohort (Table 1b & Figure 3d).

Smaller right amygdalar volumes (*p=0.035*) were noted in ADNI participants but did not meet Bonferroni significance criteria of p<0.025. Both ADNI and IMAS cohorts demonstrated similar but non-significant trends of smaller amygdalar volumes associated with elevated cortisol (Tables 1a, 1b, 1c & Figures 2, 3).

The lack of significance in IMAS is potentially attributable to inadequate power owing a much smaller sample size. An additional *post hoc* mega-analysis (Table 1c)was performed after combining ADNI and IMAS participants, which revealed elevated cortisol levels to be significantly associated with larger ventricular volumes and smaller hippocampal volumes, both noted in the right hemisphere.

## 4. Discussion

We found a significant association between elevated cortisol and smaller regional brain volumes, in an asymmetric pattern, with significant associations detected predominantly in the right hemisphere. Specifically, we demonstrated associations between elevated cortisol levels and larger lateral ventricular volumes and smaller hippocampal volumes, with similar but non-significant trends in the smaller IMAS cohort.

### 4.1 Ventricular volumes

In line with our hypothesis, increasing plasma cortisol was associated with significantly larger ventricular volumes, predominantly in the right hemisphere, in ADNI with similar non-significant trends in IMAS data. This may reflect underlying decreased periventricular white matter volumes and generalized atrophy, as can be expected based on prior studies in hypercortisolemic neuropsychiatric disorders such as bipolar disorder and schizophrenia. For example, salivary cortisol is known to affect periventricular white matter structural integrity in healthy controls and euthymic and bipolar patients (Macritchie et al., 2013) and larger ventricular volumes have been noted in patients with schizophrenia (Johnstone et al., 1976; van Erp et al., 2016).

### 4.2 Hippocampal structural change - atrophy versus re-modeling

We found significantly smaller hippocampal volumes associated with elevated cortisol, as expected based on prior literature on cortisol or hypercortisolemic neuropsychiatric conditions. Hippocampi demonstrate increased glucocorticoid receptor expression in major depressive disorder (Wang et al., 2012). Although the animal literature supports glucocorticoid-mediated neurotoxicity and apoptosis (Sapolsky, 1996), the idea of attributing cortisol-associated brain volume loss to permanent neuronal death has been called into question by human neuropathological studies, which favor the idea that brain structural changes associated with abnormally high cortisol may represent a combination of shrinkage of the neuronal soma, loss of dendritic branching, decreased adult neurogenesis, promotion of oligodendrogenesis and not necessarily neuronal cell death (Chetty et al., 2014; Curtis et al., 2007; McEwen, 2012; Swaab et al., 2005). Additionally, lower hippocampal volumes but higher neuronal density, suggestive of shrinkage and changes in neuropil, has been noted in major depression (Cobb et al., 2013).

### 4.3 Amygdalar changes

The amygdalar associations did not pass the Bonferroni significance cut off p-value < 0.025, However, we noticed with similar trends in both ADNI and IMAS cohorts, with smaller amygdalar volumes associated with elevated cortisol. Animal literature typically shows the amygdala to be a very plastic structure (Carrillo et al., 2007), with volumetric associations inconsistently noted in stress and hypercortisolemia-associated neuropsychiatric conditions, such as depression. For example, increased spine density of the basolateral amygdala was found in rats with stress (Mitra et al., 2005) and larger volumes of centromedial amygdala were noted in progressive major depressive disorder (Malykhin et al., 2018). On the other hand, smaller amygdala were seen in women with severe depression (Mervaala et al., 2000), unipolar depression (Kronenberg et al., 2009), major depressive disorder (Schmaal et al., 2016) and after chronic corticosteroid therapy (Brown et al., 2008).

### 4.4 Strengths and limitations

Strengths of our study include: (i) two well-described cohorts of older adult participants with a broad range of cognitive functioning, (ii) well-validated brain volume quantification methods for regional brain volumes (http://surfer.nmr.mgh.harvard.edu/), and (iii) adjustments for cognitive diagnosis allowing us to identify brain structural differences beyond those that translated into a cognitive deficit.

Limitations of the study include: (i) inclusion of only Caucasian participants, limiting generalizability to other ethnic groups, (ii) correlational nature of the cortisol study, as the causal direction behind HPA axis activation and brain volume loss cannot be determined clearly and is beyond the scope of this study, (iii) plasma cortisol was measured as part of a multiplex proteomic study, which may affect the sensitivity and variance of cortisol measurements, (iv) lack of a salivary cortisol measure, which has been advocated to avoid venipuncture associated stress effects and corticosteroid binding globulin associated variations on plasma cortisol measures (Gallagher et al., 2006).

Unfortunately, we were limited by the data available in ADNI and IMAS, which have a somewhat broad focus on discovering novel biomarkers for AD, and as such, collected a large variety of health measures, making it impractical to adhere to the strict standard sampling protocols used in some studies that focus specifically on HPA. Participants may have experienced some anxiety associated with undergoing MRI scanning and blood sample collection through venipuncture, and this anxiety may increase cortisol levels (Tessner et al., 2006). Even so, such an effect, if present, is unlikely to cause any systematic association of cortisol levels with brain volumes across a large sample. Due to these factors, the real strength of the association between cortisol and lower brain volume may have been underestimated in our study.

### 4.5 Future Potential

We used well-established imaging markers for stress and neuropsychiatric disorders, such as hippocampal and ventricular volumes. We would like to extend the analysis to explore every single voxel in the brain using a more unbiased approach and test the cortisol associations in both a cross-sectional and longitudinal pattern in the future.

Smaller cerebral volumes associated with elevated cortisol and stress have been shown to be reversible after correction of the hypercortisolism in Cushing’s syndrome (Bourdeau et al., 2002; Starkman et al., 1992) and after long-term recovery in anorexia nervosa (Wagner et al., 2006), suggestive of neuronal plasticity, survival and recovery after cortisol level normalization. Our study therefore encourages development of non-invasive stress and cortisol reduction strategies including meditation to reduce its detrimental effects on brain (Cahn and Polich, 2006).

Yoga and long-term meditation have demonstrated increased and volumes for the right cerebral cortex (Lazar et al., 2005), implicating its ability to potentially slow cognitive decline and degeneration. Prior animal studies have shown chronic antidepressant use to increase neurogenesis (Malberg et al., 2000), providing a strong impetus for future research involving therapeutic strategies for neurogenesis to counter the brain volume loss, such as progenitor cell transplantation, transcription factor therapy and viral implantation (Curtis et al., 2007; Martignoni et al., 1990).

## 5. Conclusion

Our study offers a useful contribution to the literature by offering further support to the theory of cortisol mediated alterations in brain parenchymal volumes. We demonstrated higher plasma cortisol was associated with larger ventricular and smaller hippocampal volumes, predominantly in the right hemisphere, regardless of age, sex or cognitive status. These brain volume changes could affect normal brain function including cognition and affect quality of life in people affected with various neuropsychiatric conditions. Our findings 1) encourage development of effective stress and cortisol reduction strategies to decrease its detrimental effects on brain and 2) offer non-invasive imaging biomarkers to test efficacy of such future therapeutic clinical trials aiming to halt or reverse the progress of volume alterations.

## Funding

**PR** was supported by Radiological Society of North America Resident Research Grant, 2017-2018. **KN** is supported by NLM R01 LM012535, R03 AG063250, and NIA R03 AG054936. NJ is additionally supported by R01AG059874. **AJS** is supported by NIA P30 AG010133, NIA K01 AG049050, NIA R01 AG061788, and NIA R01 AG19771. **PMT** is supported by U54 EB020403, P41 EB015922, P01 AG026572, P01 AG055367, RF1 AG051710, R01 AG060610, and R56 AG058854 to the ENIGMA World Aging Center.

Data used in preparation of this article were obtained from the Alzheimer’s Disease Neuroimaging Initiative (ADNI) database (adni.loni.usc.edu). As such, the investigators within the ADNI contributed to the design and implementation of ADNI and/or provided data but did not participate in analysis or writing of this report. A complete listing of ADNI investigators can be found at: http://adni.loni.usc.edu/wp-content/uploads/how_to_apply/ADNI_Acknowledgement_List.pdf.

